# Usefulness of Current sgRNA Design Guidelines and *in vitro* Cleavage Assays for Plant CRISPR/Cas Genome Editing: A Case in Eggplant (*Solanum melongena* L.)

**DOI:** 10.1101/2023.03.19.532877

**Authors:** Mark Gabriel S. Sagarbarria, John Albert M. Caraan

## Abstract

The advent of genome editing platforms such as the CRISPR/Cas9 system ushers an unprecedented speed on the development of new crop varieties that can withstand agricultural challenges of the 21^st^ century. The CRISPR/Cas9 system depends on the specificity of engineered single guide RNAs (sgRNAs). However, sgRNA design in plants can be challenging due to a multitude of design tools to choose from, many of which use guidelines that are based on animal experiments yet allow the use of plant genomes. Upon choosing sgRNAs, it is also unclear whether an *in vitro* assay is needed to validate the targeting efficiency of a particular sgRNA prior to *in vivo* delivery of the CRISPR/Cas9 system. Here, we demonstrate the *in vitro* and *in vivo* activity of four different sgRNAs that we selected based on their ability to target multiple members of the eggplant polyphenol oxidase gene family. Some sgRNAs that have high *in vitro* cleavage activity did not produce edits *in vivo*, suggesting that an *in vitro* assay may not be a reliable basis to predict sgRNAs with highly efficient *in vivo* cleavage activity. Further analysis of our sgRNAs using other design algorithms suggest that plant-validated criteria such as the presence of necessary secondary structures and appropriate base-pairing may be the reason for the discrepancy between our observed *in vitro* and *in vivo* cleavage efficiencies. However, recent reports and our data suggests that there is no guaranteed way to ensure *in vivo* cleavage of chosen sgRNAs.

**Key Message:**

- *in vitro* cleavage assay of sgRNAs was able to identify low activity sgRNAs but did not 13 reliably predict *in vivo* mutagenesis.
- Using multiple sgRNAs that meet the plant-validated parameters and have high activity *in vitro* in plant genome editing is critical to ensure success.

## INTRODUCTION

The CRISPR/Cas system has revolutionized the field of plant biology and plant breeding in recent years for its ability to produce precise and targeted induction of mutations at any region on the genome. Despite the development of other genome editing tools such as Zinc Finger Nucleases (ZFNs) or Transcription Activator-Like Effect Nuclease (TALENs), the CRISPR/Cas system has been widely adopted by many researchers as it allows for rapid generation of gene manipulation events in a simple, efficient, and specific manner with minimal to no off-target effects (Gudeta 2019). The technology has progressed to even allow chromosomal deletions (Ordon et al. 2016) and epigenetic changes (Papikian et al. 2019). This has led to the elucidation of the functions of numerous genetic elements, which has been applied for crop improvement, examples for which have been reviewed extensively (Schindele et al. 2020; Zhang et al. 2021). Genome edits are brought about through the action of the CRISPR/Cas nucleases, which must recognize a sequence downstream of the target, known as the protospacer-adjacent motif (PAM), which is 5’-NGG-3’ for *Streptococcus pyogenes* Cas9 (SpCas9), a commonly used Cas nuclease in plants. However, targeting specificity is determined by the single guide RNA or sgRNA, which usually consists of a 20-bp target-specific CRISPR RNA (crRNA) and the trans-activating crRNA (tracrRNA) which provides the necessary secondary structure for the formation of the DNA-sgRNA-Cas9 complex and activation of Cas9 (Jinek et al. 2012). Provided that Cas9 produces a double-stranded break (DSB) three bases upstream of the PAM site, a 20-bp region upstream of any PAM within the gene of interest can designated as the ‘crRNA’ to guide Cas9 to the target site. Afterwards, the crRNA can be cloned in appropriate vectors (ie., in a Cas9/sgRNA expression vector) for plant transformation.

Theoretically, there are as many target sites in a single gene as there are the number of PAM sequences. One of the many challenges is choosing which sgRNA will efficiently direct Cas9 to generate double-stranded breaks (DSBs), and eventually produce the desired edits via the cell’s error-prone repair mechanisms. To aid in this, numerous algorithms have been developed to predict the on-target efficiency of each sgRNA. However, effectiveness of these tools for plant gene editing has been questioned recently (Naim et al. 2020). On the other hand, some have recommended the use of an *in vitro* assay prior to delivery of sgRNAs *in vivo* (Holalu et al. 2019; Karmakar et al. 2021). However, this itself does not guarantee *in vivo* cleavage due to numerous factors such as chromatin accessibility (Campenhout et al. 2019). As such, there is currently no definitive protocol for sgRNA design in plants. Hence, we report our findings for sgRNA efficiency to aid future genome editing studies in lesser researched crops such as eggplant. In this study, we designed sgRNAs based on their ability to target multiple members of the polyphenol oxidase gene family in eggplant (*SmelPPO*). We then assessed the activity of each sgRNA *in vitro* prior to stable transformation and agroinfiltration. Afterwards, we compared the on-target efficiency scores of each sgRNA using different online tools, as well as design guidelines of sgRNAs validated for plants. Off-target mutations are less of a concern in plants as undesired mutations can be eliminated through backcrossing and selection during the breeding process (Tang et al. 2019). Hence, this study focused on the on-target activity of each sgRNA.

## MATERIALS AND METHODS

### Design of sgRNAs

The sgRNAs were initially designed with the aid of the online web tool Benchling (www.benchling.com), which uses the guide algorithm from Doench et al. (2016). The eggplant polyphenol oxidase 5 gene (*SmelPPO5/*SMEL4.1_08g017350.1.01) obtained from eggplant genome V4.1 (Barchi et al. 2021) was used as the input template for sgRNA design. A major consideration in choosing sgRNAs was complementation to other eggplant PPO genes, particularly in the conserved regions of the gene family (Shetty et al. 2011). Each sgRNA (from the highest on-target score) was manually aligned to *SmelPPO 4* and *6* (SMEL4.1_08g017940.1.01 and SMEL4.1_08g017330.1.01) to determine multiple target compatibility. Multiple targeting capability of each sgRNA was validated and off-targets were determined using the CasOT software (Xiao et al. 2014). Each sgRNA chosen was able to target at least two out of the three target eggplant PPO genes (*SmelPPO 4 – 6)*. We had sequenced *SmelPPO4 – 6* in our target genotype (*Solanum melongena* L. ‘PHL 11424’) to confirm sequence similarity with the reference genome. In addition, we also screened the sgRNAs to avoid internal interactions that may interfere with cleavage such as hairpins (Thyme et al. 2016).

### In vitro CRISPR/Cas9 Cleavage Assay

DNA fragments containing the target site were amplified, purified, and eluted in Tris buffer (see **Supplementary Table 1** for the list of all primers used in this study). A ribonucleoprotein reaction mix composed of synthesized sgRNAs (Macrogen, Korea; 250 ng), purified PCR products (150 ng) and Cas9 protein (500 ng) (Macrogen, Korea) in 1X NEBuffer 3.1 (100 mM NaCl, 50 mM Tris-HCl, 10 mM MgCl_2_, 100 µg/ml BSA, pH 7.9) was prepared following a minimum of 15:15:1 Cas9:sgRNA:DNA template molar reaction ratio (**Supplementary Table 2**). The reaction mixtures for each sgRNA were incubated at room temperature for 16 hours to allow the formation of the CRISPR/Cas9 ribonucleoprotein (RNP) and to achieve maximum cleavage activities. Subsequently, RNAse (10 ng) and Proteinase K (20 μg) were added into the reaction mix and incubated at 37°C for 15 minutes. The digestion products were separated in a 3.5% agarose TBE gel (1.25% Synergel gel clarifier + 1.0% agarose) and were analyzed using Gel Analyzer v. 19.1 (Lazar and Lazar 2012). The cleavage efficiencies were assessed by measuring the amount of digested products over the total amount of the initial DNA used for the *in vitro* assay.

### Preparation of CRISPR/Cas9 constructs

sgRNAs were expressed under the control of the *Arabidopsis thaliana* U6 promoter. Annealed oligos of each sgRNA were cloned into pUC19::AtU6-tracrRNA via digestion at the *Bbs*I sites. The sgRNA expression cassette was ligated into Cas9 expression vector pZH_PubiMMCas9 (Mikami et al. 2015) after digestion of both vectors with *Asc*I and *Pac*I to generate the final Cas9/sgRNA all-in-one vector. Plasmids were sequenced to verify the insertion of each sgRNA. Each Cas9/sgRNA all-in-one vector was transformed into *Agrobacterium tumefaciens* strain ‘LBA4404’.

### Delivery of CRISPR/Cas9 via *Agrobacterium*-mediated transformation and Agroinfiltration

*Agrobacterium*-mediated transformation of eggplant (*Solanum melongena* L. ‘PH 11424’) and rgeneration was carried out according to the protocol described in Sagarbarria et al. (unpublished, under review).

For agroinfiltration, an overnight culture of *A. tumefaciens* LBA4404 harboring the Cas9/sgRNA vectors was resuspended in ½ strength MS media with 1.5% sucrose and 100 μM acetosyringone. Volume was adjusted to reach an OD_600_ equal to 0.5. The *Agrobacterium* solution was resuspended in agroinfiltration buffer (½ strength MS, 1.5% sucrose and 100 mg/L acetosyringone). A 1 mL needleless syringe was used to carefully inject the abaxial surface of the third fully expanded leaf of the 4-week-old eggplants. Infiltrated portions of the leaves were marked with a permanent marker. The plants were kept in dim light conditions at 25°C under 16:8 photoperiod for 3 days before DNA extraction of the infiltrated areas.

### Mutation identification by T7E1 cleavage and heteroduplex mobility assays

Genomic DNA was extracted from acclimatized T_0_ plants, from approximately 200 mg of leaf discs (Paterson et al. 1993). The DNA pellet was resuspended in 40 – 50 μL of TE buffer and stored at -20°C until use. DNA fragments containing the target sites were amplified and purified before subjection to the T7 endonuclease I (T7E1) assay and heteroduplex mobility assay (Zhu et al. 2014). For the T7E1 cleavage assay, the protocol provided by New England Biolabs was followed. Purified PCR products amplified from T_0_ plants and infiltrated leaf samples were denatured and annealed in NEBuffer 2 with purified PCR products from wild-type samples. A total of 75 - 150 ng of hybridized PCR products were digested with T7 endonuclease (New England Biolabs, Catalogue # M0302S) for 30 minutes at 37°C. Digested products were visualized in 2% agarose gel in 1X TBE after 30 minutes of electrophoresis at 100 V. Gels were placed in 0.5% ethidium bromide solution for 10 minutes before visualization.

For the heteroduplex mobility assay (HMA), PCR products were denatured and annealed then resolved by electrophoresis in non-denaturing polyacrylamide gels containing 5% acrylamide-bisacrylamide (40% acrylamide/bis-acrylamide, 37.5:1), 1X TBE, ammonium persulfate, and TEMED. After 75 minutes of electrophoresis at 100 V, gels were placed in 0.5% ethidium bromide solution for 10 minutes before visualization.

## RESULTS

### Testing of sgRNA activity through an *in vitro* cleavage assay

Four (4) sgRNAs were chosen that could target multiple *SmelPPO* genes (**Table 1 and Figure 1)**. Synthetic sgRNAs were used with Cas9 to form the ribonucleoprotein complexes tested for *in vitro* cleavage efficiencies. Due to similar sizes of the expected cleavage products (**Supplementary Table 3**), we took the amount for all observed digested bands in computation of the cleavage efficiency (**Supplementary Table 4**).

**Table 1.**
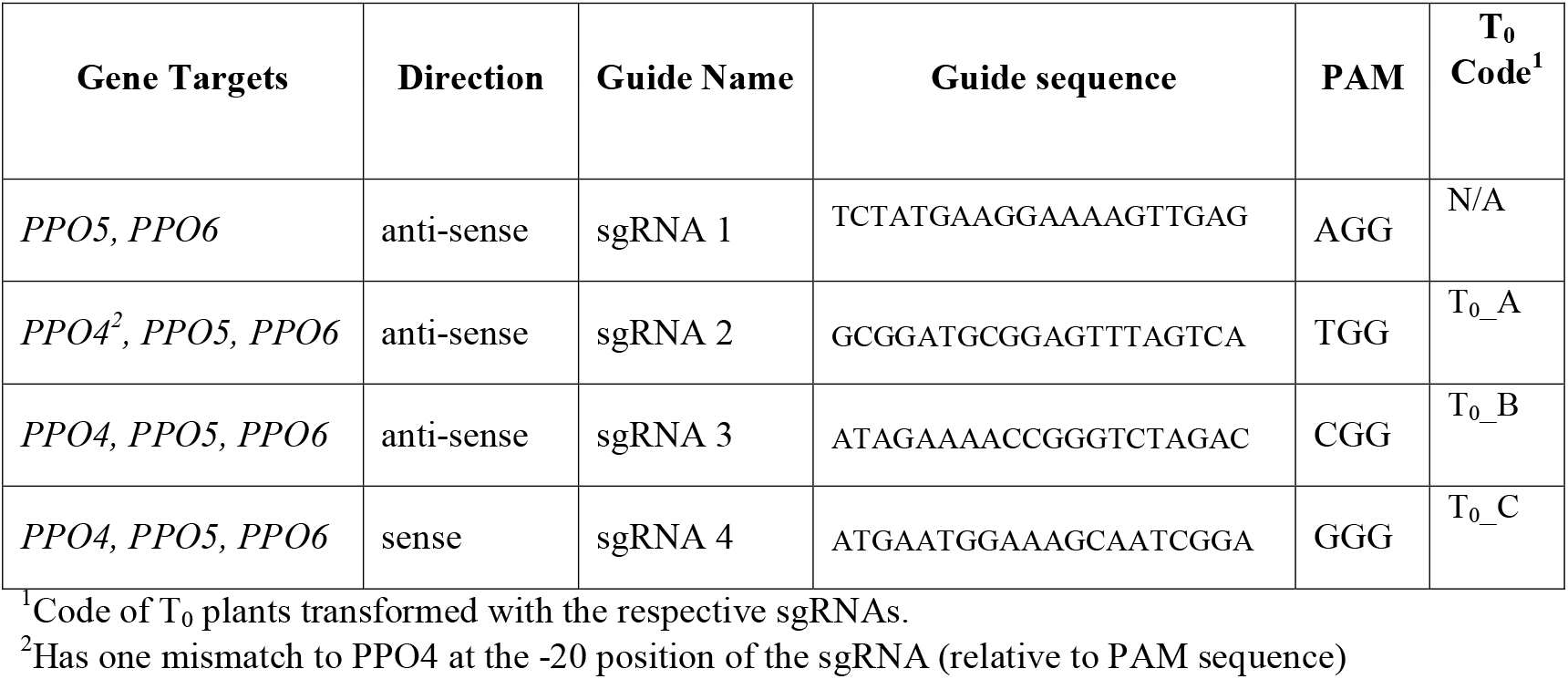
List of sgRNAs that could target multiple *SmelPPO* genes.

**Figure 1.**
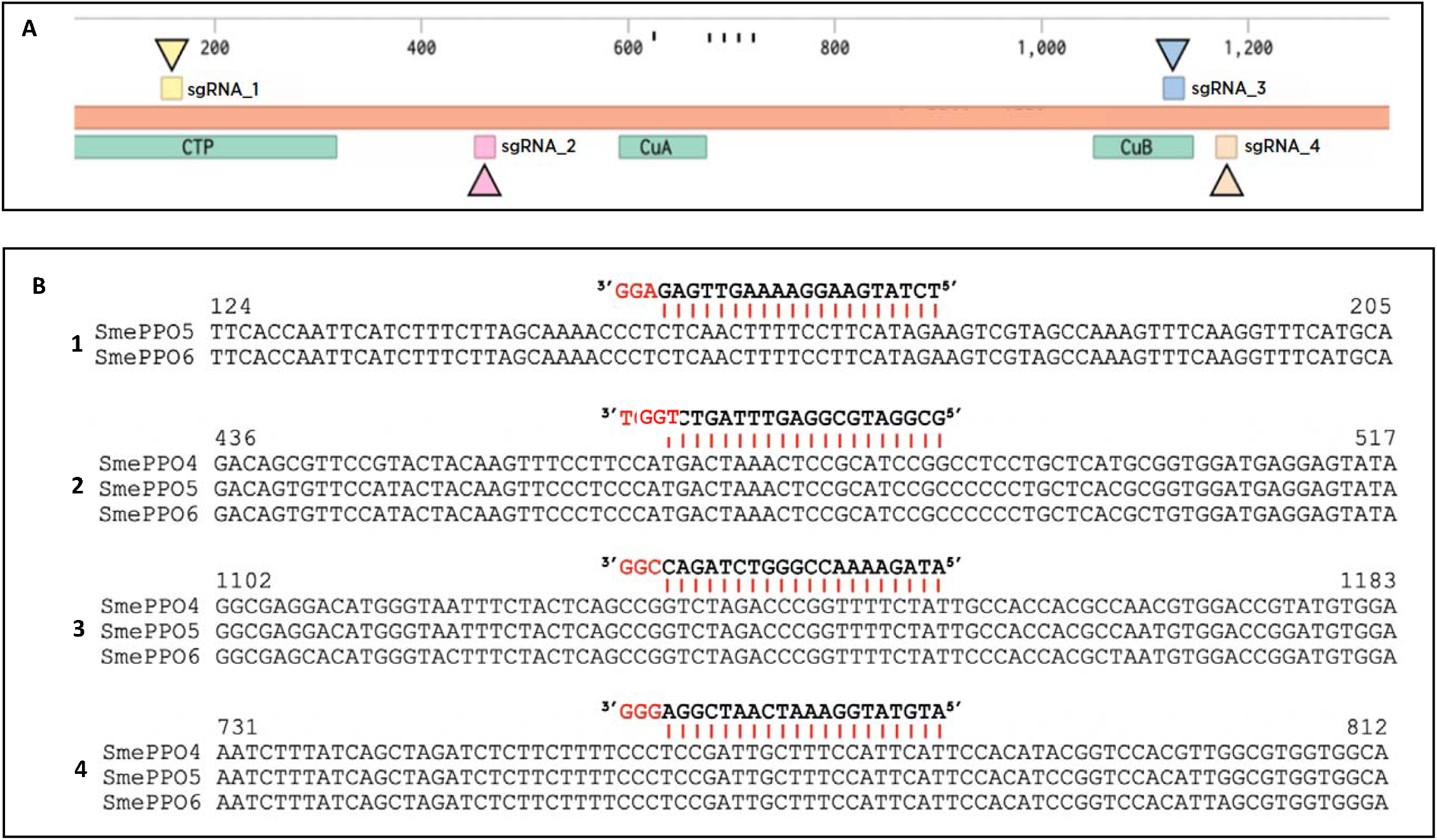
**(A)** Linear sequence map of *SmelPPO5* showing the target sites of the designed sgRNAs at various gene domains. Target sites of the sgRNAs were labeled as follows: Yellow = Chloroplast Transit Peptide Domain (CTP); Pink = Near the Copper A Binding Domain (CuA); Blue and Cream = Copper B Binding Domain (CuB). **(B)** sgRNA target sites in multiple *SmelPPO* genes (1) sgRNA_1 targeting *SmelPPO*5 and 6, (2) sgRNA_2 targeting *SmelPPO* 4, 5, and 6, (3) sgRNA_3targeting *SmelPPO* 4, 5, and 6, and (4) sgRNA_4 targeting *SmelPPO* 4, 5, and 6

From the observed digestion products of the RNP cleavage assay, all four (4) sgRNAs were able to direct the Cas9 endonuclease to the target sites to perform DSBs, albeit at different efficiencies (**Figure 2**). sgRNA_1 targeting the CTP domain was able to cleave its target genes (*SmelPPO5* and *6*) at an *in vitro* efficiency rate of 85.15% (+++). sgRNA_2 targeting near the CuA binding domain, on the other hand, obtained highly efficient *in vitro* cleavage activities on *SmelPPO5* and *6* at a rate of 84.70% (+++), but no cleavage activity was observed in *SmelPPO4*. sgRNA_2 has a single mismatch (guanine-guanine) at 20^th^ base upstream of the PAM for *SmelPPO4*. Initially, we had expected that this mismatch would be tolerated by the CRISPR/Cas9 system, as it has been reported that sgRNAs can exhibit function when the mismatches occur at the most distal end from the PAM (Anderson et al. 2015). However, since other targets are present that have perfect complementation, *SmelPPO5* and *6* may have been preferred for cleavage. Meanwhile, sgRNA_3 targeting the CuB binding domain generated the least efficient *in vitro* cleavage scores at a rate of 44.40% (+) among the four sgRNAs. This may be because of nucleotide preferences of Cas9 particularly at the seed sequence (Doench et. al., 2014). At the 1^st^ base upstream of the PAM (position 20 relative to the target gene), Cas9 favors a guanine residue the most while greatly disfavors a cytosine residue. In case of sgRNA_3, the residue which immediately follows the PAM recognition sequence is cytosine. Lastly, we observed that sgRNA_4 was able to produce highest *in vitro* efficiency score at a rate of 87.48% (+++) across all target genes *SmelPPO genes* (4, 5, and 6). Since all the tested sgRNAs had observed cleavage *in vitro*, we forwarded all sgRNAs in transformation experiments.

**Figure 2.**
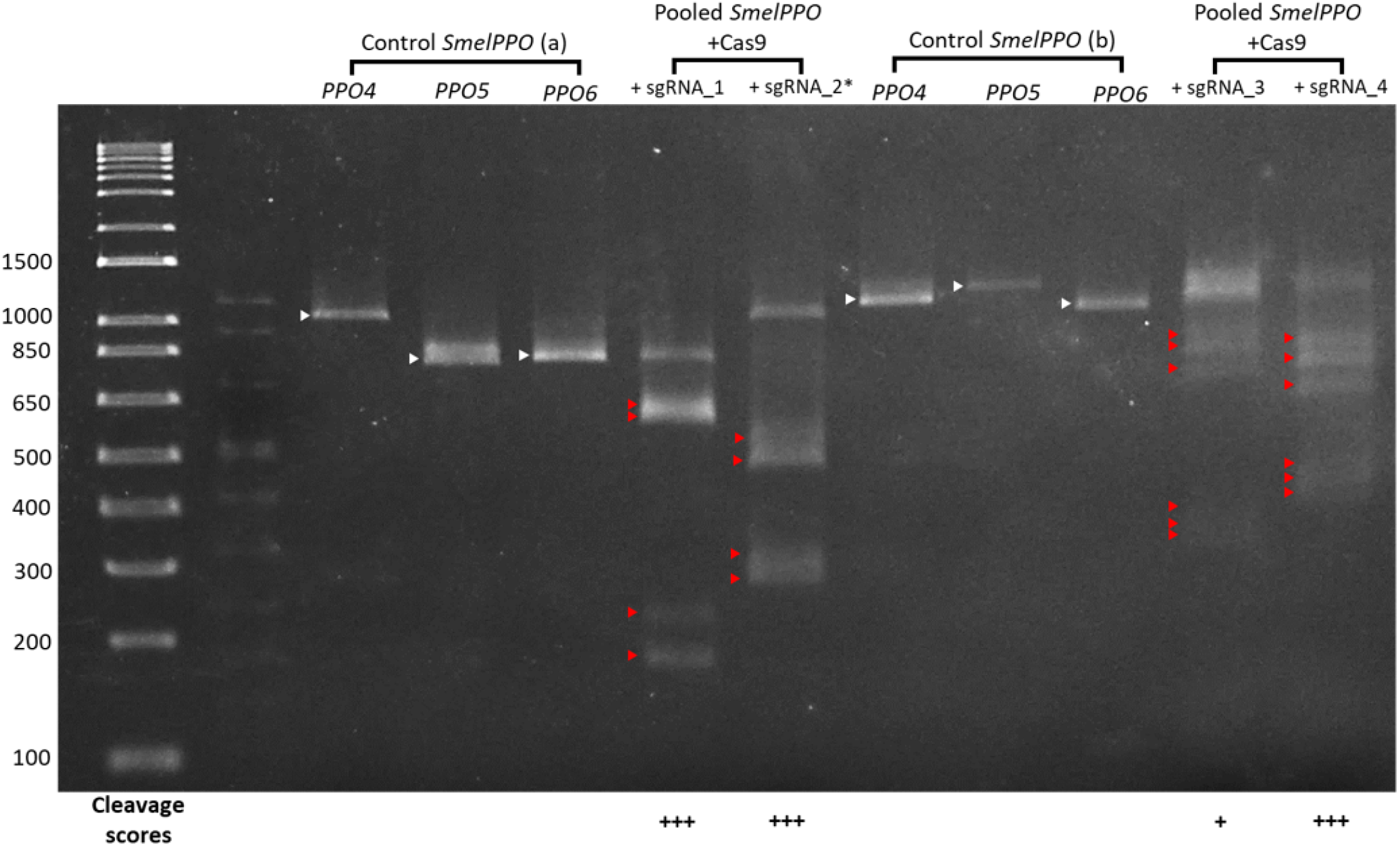
*In vitro* cleavage assay of the four (4) sgRNA/Cas9 ribonucleoprotein complex on the pooled PCR-amplified *Smel*PPO genes from Philippine Eggplant cv. ‘Mistisa’ (PH11424). (White arrows) Wild type *SmelPPO* (*4, 5, and* 6)DNA used as control. (Red arrows) Digested *SmelPPO* (*4, 5, and/or* 6) DNA after 16 hours of cleavage assay. Cleavage scores: (+) ≤60% cleavage; (++) 61-80% cleavage; (+++) ≥81% cleavage. \**In vitro* cleavage activity only observed for *SmelPPO5* and *SmelPPO6*

### Editing the eggplant PPO gene family through *Agrobacterium*-mediated stable transformation and agroinfiltration

The construct, pZH_PubiMMCas9 (**Figure 3A**), which encodes a Cas9 protein under control of the maize ubiquitin promoter and an sgRNA expression cassette under the control of the *A. thaliana* U6 promoter, was used in transformation of eggplant cotyledon explants. A total of 9 plants were regenerated and acclimatized across sgRNA_2 - 4. All obtained regenerants tested positive for the presence of *Cas9* and *HPT* transgenes (**Supplementary Figure 1**), indicating successful transformation. The T7 endonuclease I (T7E1) assay and heteroduplex mobility assays (HMA) were used to detect mutations in all the target sites for each sgRNA. Mutations were only detected from plants transformed with sgRNA_4/T_0__C plants (**Figure 3C-D, Figure 3F-G, Figure 4A-B**). For plants transformed with sgRNA_2/T_0__A plants, multiple bands for each gene target were detected in the HMA even in control samples (**Figure 3D**). However, no cleavage products were observed in the T7E1 assay (**Figure 3C**). We attribute these to be ‘shadow’ bands that formed due to the different structural formations that may have formed in the PCR products that were separated in non-denaturing PAGE. Since these bands were also present on the control samples, but mismatch cleavage was not observed in the T7E1 assay, the multiple observed bands are most likely the same DNA fragment in different structural conformations. We had also observed that samples positive in the T7E1 assay were also positive in the HMA aside from T_0__C2 for *SmelPPO6*, which tested positive in the HMA but negative in the T7E1 assay. This could be because the T7E1 assay is considered less sensitive in accurately reporting the percentage of edited cells (Sentmanat et al. 2018), but is still useful as it does not overestimate or produce false positive results (Bell et al. 2014).

**Figure 3.**
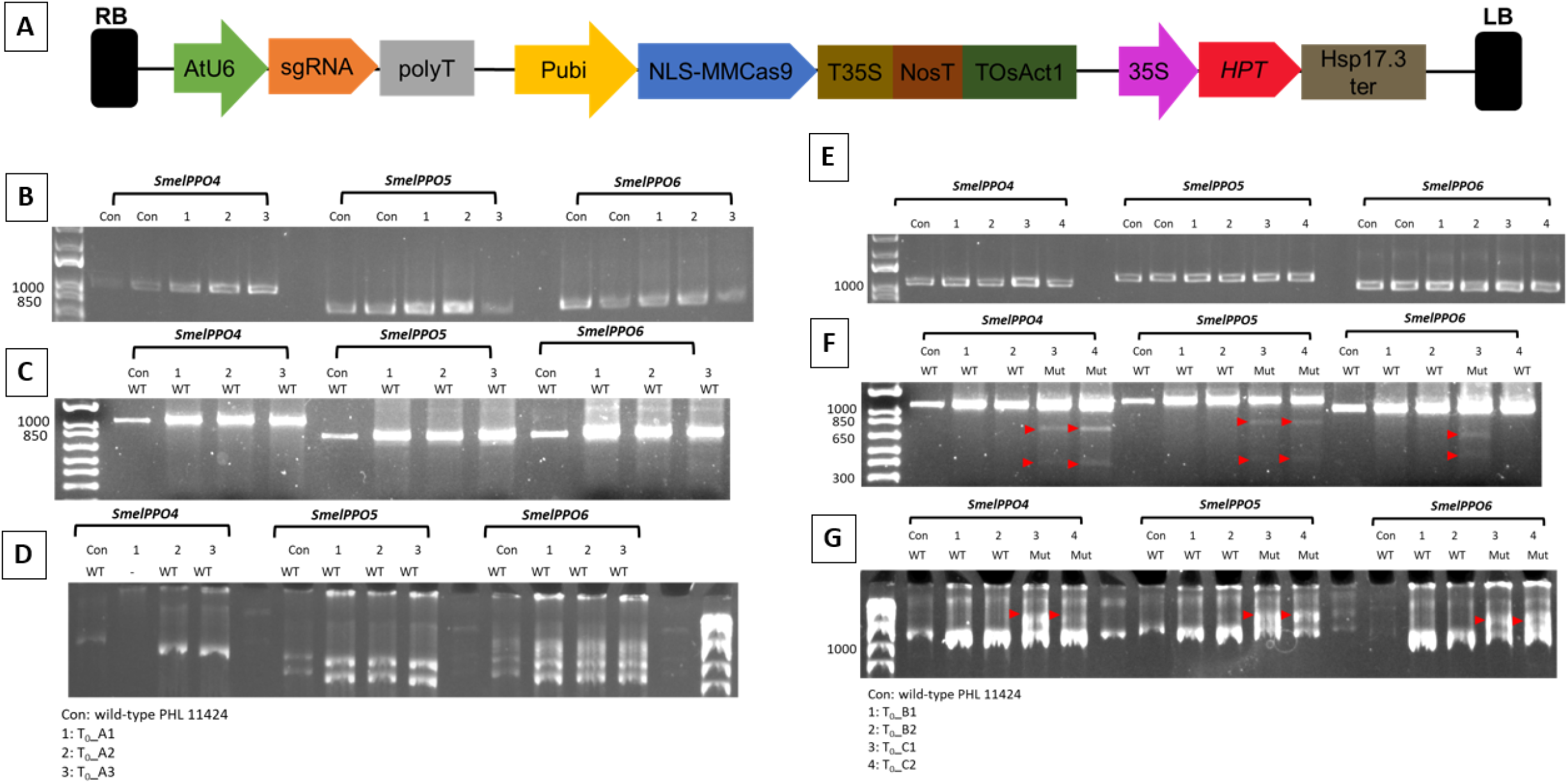
**(A)** Cas9/sgRNA all-in-one vector plasmid based on pZH_PubiMMCas9. **(B)** Purified PCR products of *SmelPPO4-6* in T0_A plants. **(C)** T7E1 assay of PCR products of *SmelPPO4-6* in T0_A plants hybridized with PCR products from wild-type plants in 2% agarose gel. **(D)** HMA of PCR products of of *SmelPPO4-6* in T0_A plants in 5% PAGE. **(E)** Purified PCR products of SmelPPO4-6 in T0_B and T0_C plants. **(F)** T7E1 assay of PCR products of *SmelPPO4-6* in T0_B and T0_C plants hybridized with PCR products from wild-type plants in 2% agarose gel. **(G)** HMA of PCR products of of *SmelPPO4-6* in T0_B and T0_C plants in 5% PAGE. Red arrows indicate presence of cleavage products for T7E1 assay and heteroduplexes from HMA.

**Figure 4.**
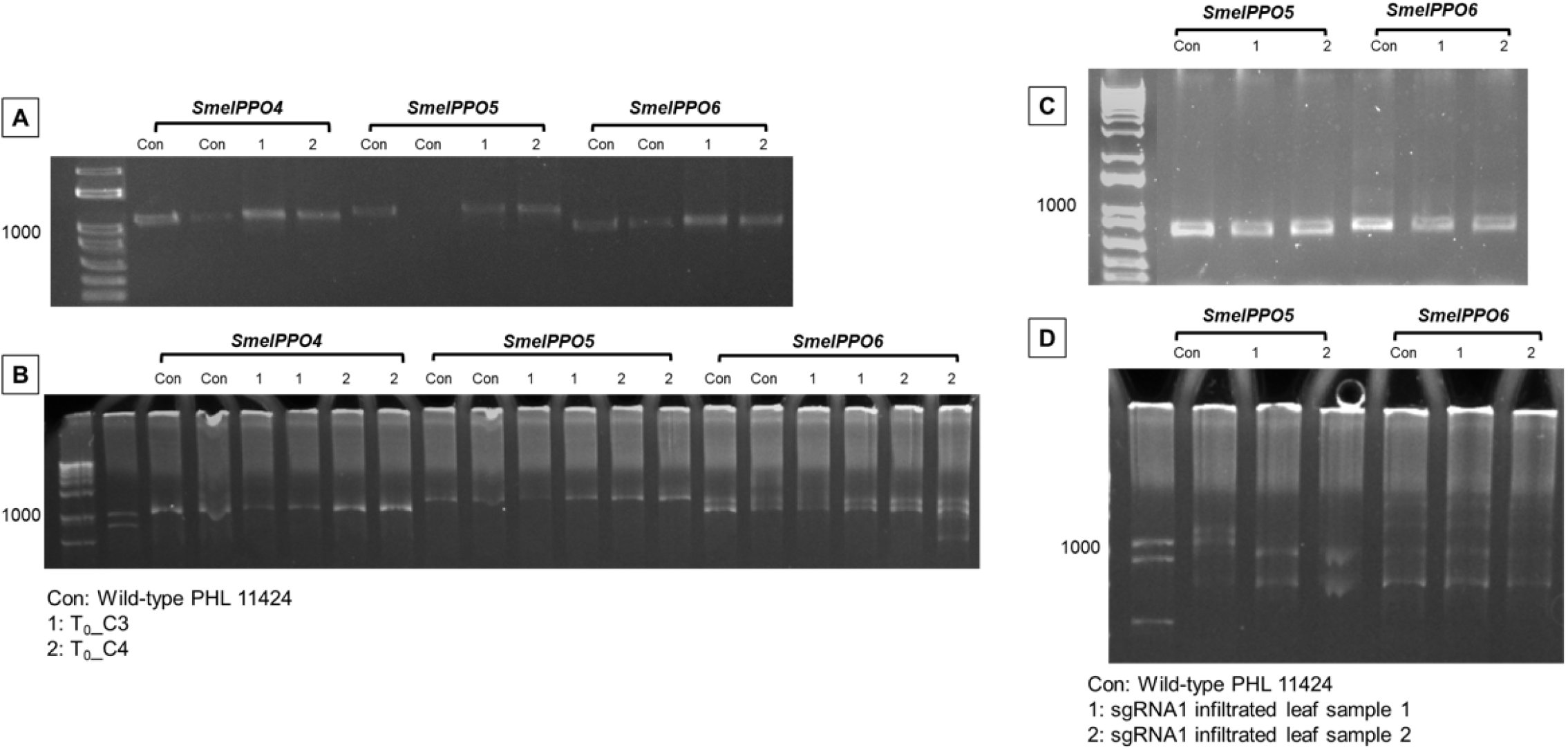
**(A)** T7E1 assay of PCR products of *SmelPPO4-6* in T0_C plants hybridized with PCR products from wild-type plants in 2% agarose gel. **(B)** HMA of PCR products of of *SmelPPO4-6* in T0_C plants in 5% PAGE. (C) T7E1 assay of PCR products of *SmelPPO5-6* from sgRNA1 agroinfiltrated leaves hybridized with PCR products from wild-type plants in 2% agarose gel. (D) HMA of PCR products of of *SmelPPO5-6* from sgRNA1 agroinfiltrated leaves in 5% PAGE.

Unfortunately, no regenerants were obtained for sgRNA_1. We included a transient transformation of sgRNA1 into eggplant leaves via agroinfiltration to determine if sgRNA_1 could produce edits *in vivo*. However, no edits were detected in both T7E1 and HMA (**Figure 4C-D**).

### Comparison of *in vitro* and *in vivo* genome editing using secondary structure guidelines and sgRNA design scores

In order to understand the potential reasons that could have caused the discrepancy between the *in vitro* and *in vivo* cleavage/mutagenesis, we had investigated the predicted efficiency of each sgRNA across numerous design platforms and the secondary structure of each sgRNA. Five total tools were used to predict the efficiency of the chosen sgRNAs: 1) CRISPR-P (Liu et al. 2017), 2) CRISPRDB (Chen and Wang 2022), 3) CRISPOR (Concordet and Haeussler 2018), 4) CCTop (Stemmer et al. 2015), 5) CasDesigner (Park et al. 2015). The Doench et al. (2016) score was used for CRISPOR, since according to the CRISPOR manual, this performed better for U6 datasets. It is important to note that only CRISPOR supported the use of the eggplant reference genome to check for off-targets. Thus, only on-target scores were obtained from these platforms. Lastly, secondary structures and internal base-pairing of each sgRNA was checked using Mfold (www.unafold.org) (Zuker 2003), using guidelines from highly efficient sgRNAs in plants (Liang et al. 2016).

We also investigated the major characteristics of an effective sgRNA (Doench et al. 2016; Mohammadhassan et al. 2022), which are 1) number of targets/off-targets, 2) %GC content, 3) sgRNA length, 4) sgRNA sequence, and 5) secondary structure. For the off-targets, it is assumed that the same sgRNA across all targets and off-target sites would have competition for binding. However, all the tested sgRNAs have a high number of mismatches for potential off-targets, with sgRNA_4 being the lowest at two mismatches for one off-target site (**Table 2**). The CRISPR/Cas9 system has low tolerance for mismatches, with less than 5% of crRNAs being functional at sites with two mismatches (Anderson et al. 2015). Using the same assumption for number of targets, sgRNA_3 and 4 would be the least efficient with sgRNA_1 having higher mutagenesis rates. However, sgRNA4 was the only sgRNA to produce mutagenesis *in vivo* despite having the most potential for an off-target site. Therefore, the number of targets/off-targets does not seem to be a major factor that would explain the observed mutagenesis in the four sgRNAs. The second factor, %GC content is within the reported optimal range of 30 – 80% (Liang et al. 2016). It would be difficult to attribute the vast difference in observed *in vitro* and *in vivo* cleavage to a difference of 5 – 10% GC content, and so this may not be a major factor in sgRNA editing efficiency. For sgRNA length, all sgRNA used are 20 nt long, which was found to be the most efficient in rice (Liu et al. 2022). Thus, this should also not be a major factor for consideration. In addition to sgRNA length, distance between the start codon and the PAM site has been reported to have a significant impact on cleavage efficiency (Matson et al. 2019). However, the distance between the non-functional sgRNA_3 and functional sgRNA_4 is on average 65 nt across their three target sites, ruling out this factor. For sgRNA sequence, animal studies have reported certain nucleotide positions attributed to more functional sgRNA (Doench et al. 2014). But for sgRNAs validated in plants, it was found that there is no nucleotide preference for functional sgRNAs (Liang et al. 2016), ruling out this factor. For sgRNA secondary structure, *in silico* prediction of the secondary structure of each sgRNA revealed that all four sgRNAs had a secondary structure that contained all the necessary stem loops (RAR, SL2, and SL3) found in validated plant sgRNAs (Liang et al. 2016). However, all sgRNAs except sgRNA_4 had excessive base-pairing between the guide sequence (crRNA) and the scaffold (tracrRNA), exceeding the maximum of 12 bases paired for sgRNA_1 – 3 and more than the maximum of 7 consecutive base pairs for sgRNA_1 (**Supplementary Table 5**). This suggests that the base pairings within sgRNA_1 – 3 are favored and affected the base pairing of the guide sequence with its target DNA, reducing mutagenesis rates *in vivo*. We compared these results with scores for online prediction tools and observed that only CRISPR-P was able to give low confidence scores to sgRNA_1 – 3 (**Table 2**), as only CRISPR-P considers these guidelines for plant validated sgRNAs.

**Table 2.** Comparison of different factors affecting sgRNA efficiency in comparison to observed *in vitro* and *in vivo* cleavage results.

| Targets | Name | GC (%) | Mismatch number<sup>1</sup> | Off-targets<sup>2</sup> | sgRNA Rankings |  |  |  |  | Liang et al. (2016) Guidelines<sup>3</sup> | *In vitro* | *In vivo* |
| --- | --- | --- | --- | --- | --- | --- | --- | --- | --- | --- | --- | --- |
|  |  |  |  |  | CRISPR-P | CRISPRDB | CRISPOR | CCTop | Cas-Designer |  |  |  |
| PPO5, PPO6 | sgRNA _1 | 35 | 3 | 5 | 0.0769 | 80.5 | 66 | 0.76 | 82 | Fail | +++ | No |
| PPO4, PPO5, PPO6 | sgRNA _2 | 55 | 4 | 4 | 0.039 | 51.7 | 46 | 0.69 | 66.6 | Fail | +++<sup>4</sup> | No |
| PPO4, PPO5, PPO6 | sgRNA _3 | 45 | 3 | 1 | 0.0086 | 30.2 | 48 | 0.69 | 62.7 | Fail | + | No |
| PPO4, PPO5, PPO6 | sgRNA _4 | 40 | 2 | 1 | 0.6809 | 92.2 | 66 | 0.49 | 71.8 | Pass | +++ | Yes |

<sup>1</sup>minimum number of mistakes between sgRNA sequences and possible off-target sequences
<sup>2</sup>Number of off-targets at that have the minimum mismatch number
<sup>3</sup>See Supplementary Table 2 for a detailed breakdown for each sgRNA
<sup>4</sup>*In vitro* cleavage activity only observed for *SmelPPO5* and *SmelPPO6*

## DISCUSSION

It has been previously suggested that sgRNA ranking tools for genome editing in plants were not useful in avoiding less cleavable sites or selecting highly susceptible sites, and that it would be more effective to choose PAM sites in coding exons for gene disruption rather than choose solely from prediction scores (Naim et al. 2020). We took a similar approach in that we prioritized sgRNAs that could target multiple members of the eggplant polyphenol oxidase (*SmelPPO*) gene family and narrowing down the list with the ones with high on-target scores from our chosen design platform, low number of off-targets, and checking for hairpin structures that could interfere with cleavage. Due to other reports that sgRNA prediction algorithms are not able to consistently predict functional sgRNAs (Gagnon et al. 2014; Thyme et al. 2016; Grainger et al. 2017), we subjected each sgRNA to an *in vitro* assay. All four of our chosen sgRNAs were able to produce cleavage *in vitro*, which we had initially assumed would result in at least some level of *in vivo* activity. Some reports had used an *in vitro* cleavage assay to screen for efficient sgRNA prior to delivery of the Cas9/sgRNA in plants (Malnoy et al. 2016; Liang et al. 2017; Bente et al. 2020). Although in zebrafish, sgRNA with at least some *in vitro* activity were found to produce mutations *in vivo*, while those with no *in vitro* activity failed to produce mutations *in vivo* (Grainger et al. 2017). However, we have observed that sgRNA with *in vitro* activity will not necessarily produce mutations *in vivo* in eggplant. We observed that the *in vitro* assay could be useful in predicting which sgRNAs will have low *in vivo* activity but could not discriminate the sgRNAs that had high *in vitro* activity but low activity *in vivo*. The results support the finding that synthetic RNA guides do not seem to have a link between efficiency and sequence at all (Haeussler and Concordet 2023).

In this case study, we have narrowed down the major factor for the discrepancy between *in vitro* and *in vivo* mutagenesis to be the presence of the needed sgRNA secondary structures, as well as the number of base pairings within the sgRNA. Still, these parameters may not be the ultimate basis for deciding on which sgRNAs to choose, as no correlation was observed with editing frequencies and sgRNA ranking scores (including CRISPR-P) (Naim et al. 2020). Activity *in vivo* can be further reduced as some genomic regions may not be susceptible to Cas9 cleavage due to limited chromatin accessibility (Uusi-Mäkelä et al. 2018). This is not considered in current sgRNA design algorithms yet. For this study, we assume that *SmelPPO 4 – 6* are available for cleavage by Cas9 as these are expressed in the plant (leaves, stems, flowers, fruits) at relatively abundant levels in comparison to house-keeping tubulin genes (Shetty et al. 2011). However, chromatin accessibility is a major determinant of Cas9 cleavage (Chung et al. 2020) and should be considered in future genome editing experiments should data of plant epigenetic landscapes become available.

What we collect from our results and recent reports on sgRNA cleavage efficiency in plants is that, while using sgRNA design and ranking tools are convenient and using an *in vitro* assay to weed-out low activity sgRNAs may be useful, there is no guaranteed way to ensure *in vivo* cleavage of the chosen sgRNA. Hence, using multiple sgRNAs that meet the plant-validated parameters and have high activity *in vitro* in plant genome editing is critical to ensure success.

## RECOMMENDATIONS

A major constraint of our study is that we had initially used Benchling for the design of our initial list of sgRNAs. The Benchling platforms still follows the Doench et al. (2016) algorithm which does not consider guidelines for plant validated sgRNAs. Perhaps if we used the CRIPSR-P platform from the start, we would have had more success in finding efficient sgRNAs. In addition, we had only used the T7E1 assay and HMA to detect mutations. Since these are PCR-based assays, it is biased towards the more abundant template and small numbers of mutations may not be detected (Zischewski et al. 2017). It could be possible that edits are present from plants transformed with sgRNA1-3 but were not detected in the T7E1 assay and HMA. Further mutation screening should be done in T_1_ generations using more accurate methods such as sequencing.

## DECLARATIONS

### Authors’ contributions

Mark Gabriel S. Sagarbarria conceptualized the study and both authors contributed to the study design. John Albert M. Caraan performed most of the *in vitro* experiments and construct assembly. Mark Gabriel S. Sagarbarria performed the T7E1 assay and HMA. Transformation, regeneration, material preparation, data collection and analysis were performed by both authors. The first draft of the manuscript was written by Mark Gabriel S. Sagarbarria and all authors commented on previous versions of the manuscript. All authors read and approved the final manuscript.

## Acknowledgements

We thank Dr. Kazuo N. Watanabe, who provided the sgRNA cloning vector and all-in-one Cas9/sgRNA vector. We thank Dr. Desiree M. Hautea for her research supervision in the early stages of this work. We also thank Ms. Rowena B. Frankie for her assistance with tissue culture and Mr. Angelo Layos for assistance in DNA preparation and PCR.

