## Supplementary Table 1 for "Usefulness of Current sgRNA Design Guidelines and *in vitro* Cleavage Assays for Plant CRISPR/Cas Genome Editing: A Case in Eggplant (*Solanum melongena* L.)"

**Supplementary Table 1.** List of all primers used in this study.

| Primer Name | Sequence (5' - 3') | Remarks |
| --- | --- | --- |
| MMCas9-F | GACCCGCAAGTCTGAAGAAACT | cas9 amplification |
| MMCas9-R | TCTTGAGGAGATCGTGGTACGT |  |
| HPT-F | CGAGAGCCTGACCTATTGCATC | HPT amplification |
| HPT-R | GAAATTGCCGTCAACCAAGCTC |  |
| sgRNA-F | TAATCTTCAAAAGGCCCCTGGG | plasmid verification |
| sgRNA-R | AGCTCCCATCCACTAGTGATCA |  |
| PPO4 (A) - F | TTTCAAGGTTTCATGCAACGCC | amplification of PPO4 with sgRNA 2 target site for in vitro and mutation identification assays |
| PPO4 (A) - R | TCCCTCCGATTGCTTTCCATTC |  |
| PPO5 (A) - F | GACTCAGCAACATGTTTCACGT | amplification of PPO5 with sgRNA 1-2 target site for in vitro and mutation identification assays |
| PPO5 (A) - R | TGGTAAAGCGAAAGTTGGATCA |  |
| PPO6 (A) - F | GCAACATGTTTCACGTTTGTGC | amplification of PPO6 with sgRNA 1-2 target site for in vitro and mutation identification assays |
| PPO6 (A) - R | TGCCTTTTGGATGGTCCCAATT |  |
| PPO4 (B) - F | AGGTTCATCCCTTTACGATGCA | amplification of PPO4 with sgRNA 3-4 target site for in vitro and mutation identification assays |
| PPO4 (B) - R | GAGCACAGTAACCAGCAATCCA |  |
| PPO5 (B) - F | TCCCATGTTTGATCGTGAAGGT | amplification of PPO5 with sgRNA 3-4 target site for in vitro and mutation identification assays |
| PPO5 (B) - R | ACGTAAAGTCAACAATGGGAAAGAG |  |
| PPO6 (B) - F | AAAAGGCATGCGTATACCTCCC | amplification of PPO6 with sgRNA 3-4 target site for in vitro and mutation identification assays |
| PPO6 (B) - R | ACGATCTCCGCATTCTCAATGG |  |
| sgRNA1-F | ATTGTCTATGAAGGAAAAGTTGAG | sgRNA cloning |
| sgRNA1-R | AAACCTCAACTTTTCCTTCATAGA |  |
| sgRNA2-F | ATTGGCGGATGCGGAGTTTAGTCA | sgRNA cloning |
| sgRNA2-R | AAACTGACTAAACTCCGCATCCGC |  |
| sgRNA3-F | ATTGATAGAAAACCGGGTCTAGAC | sgRNA cloning |
| sgRNA3-R | AAACGTCTAGACCCGGTTTTCTAT |  |
| sgRNA4-F | ATTGATGAATGGAAAGCAATCGGA | sgRNA cloning |
| sgRNA4-R | AAACTCCGATTGCTTTCCATTCAT |  |
