## Supplementary Table 2 for "Usefulness of Current sgRNA Design Guidelines and *in vitro* Cleavage Assays for Plant CRISPR/Cas Genome Editing: A Case in Eggplant (*Solanum melongena* L.)"

| **sgRNA Targets** | **DNA Template (Purified PCR Product)*** | | **Cas9**  **500 ng (uL)** | **sgRNA**  **250 ng (uL)** | **10X NEB Buffer (uL)** | **Sigma Water (to volume)**** |
| --- | --- | --- | --- | --- | --- | --- |
|  | **[ ](ng/uL)** | **150 ng (uL)** |  |  |  |  |
| **CTP** |  | **5.3** | **2** | **2** | **2** | 8.7 |
| PPO 5 (a) | 62.083 | 2.4 |  |  |  |  |
| PPO 6 (a) | 52.165 | 2.9 |  |  |  |  |
| **CuA** |  | **9.1** | **2.5** | **3** | **3** | 12.4 |
| PPO 4 (a) | 39.742 | 3.8 |  |  |  |  |
| PPO 5 (a) | 62.083 | 2.4 |  |  |  |  |
| PPO 6 (a) | 52.165 | 2.9 |  |  |  |  |
| **CuB1** |  | **15.3** | **2.5** | **3** | **3** | 6.2 |
| PPO 4 (b) | 38.652 | 3.9 |  |  |  |  |
| PPO 5 (b) | 21.71 | 6.9 |  |  |  |  |
| PPO 6 (b) | 33.09 | 4.5 |  |  |  |  |
| **CuB2** |  | **15.3** | **2.5** | **3** | **3** | 6.2 |
| PPO 4 (b) | 38.652 | 3.9 |  |  |  |  |
| PPO 5 (b) | 21.71 | 6.9 |  |  |  |  |
| PPO 6 (b) | 33.09 | 4.5 |  |  |  |  |

**Supplementary Table 2.** Reaction mixture for the *in vitro* RNP cleavage assay (minimum molar reaction of 15:15:1 Cas9:sgRNA:DNA Template).

*DNA templates of the sgRNA targets were pooled into one reaction mix
**10 uL reaction volume per sgRNA target
