## Supplementary Table 3 for "Usefulness of Current sgRNA Design Guidelines and *in vitro* Cleavage Assays for Plant CRISPR/Cas Genome Editing: A Case in Eggplant (*Solanum melongena* L.)"

**Supplementary Table 3.** Expected fragment sizes after incubation with the Cas9/sgRNA RNP for each target gene.

| **Target** | **Undigested band (bp)** | **+sgRNA1** | | **+sgRNA2** | | **+sgRNA3** | | **+sgRNA4** | |
| --- | --- | --- | --- | --- | --- | --- | --- | --- | --- |
|  |  | **Expected fragment size 1 (bp)** | **Expected fragment size 2 (bp)** | **Expected fragment size 1 (bp)** | **Expected fragment size 2 (bp)** | **Expected fragment size 1 (bp)** | **Expected fragment size 2 (bp)** | **Expected fragment size 1 (bp)** | **Expected fragment size 2 (bp)** |
| ***SmelPPO4* (a)** | 1016 | – | – | 736 | 280 | – | – | – | – |
| ***SmelPPO5* (a)** | 784 | 590 | 194 | 496 | 288 | – | – | – | – |
| ***SmelPPO5* (a)** | 800 | 619 | 181 | 484 | 316 | – | – | – | – |
| ***SmelPPO4* (b)** | 1100 | – | – | – | – | 777 | 323 | 714 | 386 |
| ***SmelPPO5* (b)** | 1199 | – | – | – | – | 857 | 342 | 765 | 404 |
| ***SmelPPO5* (b)** | 1041 | – | – | – | – | 682 | 359 | 619 | 422 |
