## Supplementary Table 4 for "Usefulness of Current sgRNA Design Guidelines and *in vitro* Cleavage Assays for Plant CRISPR/Cas Genome Editing: A Case in Eggplant (*Solanum melongena* L.)"

**Supplementary Table 4*.*** *In vitro* RNP cleavage scores obtained using Gel Analyzer v. 19.1.

| **sgRNA code** | **Target** | **Raw Volume of Bands Observed (Digested + Undigested)** | **Raw Volume of Digested DNA Products (Pooled)** | **Initial concentration of DNA used for in vitro assay (ng)** | **% Cleavage Efficiency*** | **Cleavage Score**** |
| --- | --- | --- | --- | --- | --- | --- |
| sgRNA_1 | CTP domain of *SmelPPO5* & *6* | 1138 | 969 | 300 | 85.15 | +++ |
| sgRNA_2 | Near CuA domain of *SmelPPO4, 5* & *6* | 928 | 786 | 450 | 84.70 | +++ |
| sgRNA_3 | CuB domain of *SmelPPO4, 5* & *6* | 964 | 428 | 450 | 44.40 | + |
| sgRNA_4 | Near CuB domain of *SmelPPO4, 5* & *6* | 759 | 664 | 450 | 87.48 | +++ |

*(Raw volume of digested bands/Raw volume of total bands observed)*100
**	(-) no digestion products,

(+) ≤60%

(++) 61-80%

(+++) ≥81%
