## Supplementary Table 5 for "Usefulness of Current sgRNA Design Guidelines and *in vitro* Cleavage Assays for Plant CRISPR/Cas Genome Editing: A Case in Eggplant (*Solanum melongena* L.)"

**Supplementary Table 5.** Analysis of secondary structures and base-pairing within each sgRNA using Mfold.

| Guide Name | Guide sequence | PAM | # of predicted secondary structures (Mfold) | dG | Liang et al. (2016) Guidelines | | | | Pass or Fail |
| --- | --- | --- | --- | --- | --- | --- | --- | --- | --- |
|  |  |  |  |  | Present Stem Loops^1^ | TBPs^2^ | CBPs^3^ | IBPs^4^ |  |
| sgRNA_1 | TCTATGAAGGAAAAGTTGAG | AGG | 2 | -28.7 | RAR, SL2, SL3 | 14 | 14 | 0 | Fail |
|  |  |  |  | -28 | RAR | 14 | 14 | 0 | Fail |
| sgRNA_2 | GCGGATGCGGAGTTTAGTCA | TGG | 2 | -33.6 | RAR, SL2, SL3 | 13 | 6 | 0 | Fail |
|  |  |  |  | -32.1 | RAR, SL2, SL3 | 14 | 6 | 0 | Fail |
| sgRNA_3 | ATAGAAAACCGGGTCTAGAC | CGG | 2 | -30.1 | RAR, SL2, SL3 | 13 | 5 | 0 | Fail |
|  |  |  |  | -28.8 | RAR | 13 | 6 | 0 | Fail |
| sgRNA_4 | ATGAATGGAAAGCAATCGGA | GGG | 2 | -27.1 | RAR, SL2, SL3 | 9 | 6 | 0 | Pass |
|  |  |  |  | -26.4 | RAR | 9 | 6 | 0 | Fail |

^1^RAR = repeat anti-repeat, SL1 = stem loop 1, SL2 = stem loop 2, SL3 = stem loop 3

^2^TBPs (total base pairs): total bases of the crRNA paired to the tracrRNA scaffold

^3^CBPs (consecutive base pairs): most number of consecutive base pairs of the crRNA to the tracrRNA scaffold

^4^IBPs (internal base pairs): total bases paired within the crRNA
