## Supplementary figures and images for "Usefulness of Current sgRNA Design Guidelines and *in vitro* Cleavage Assays for Plant CRISPR/Cas Genome Editing: A Case in Eggplant (*Solanum melongena* L.)"

### Supplementary Figure 1

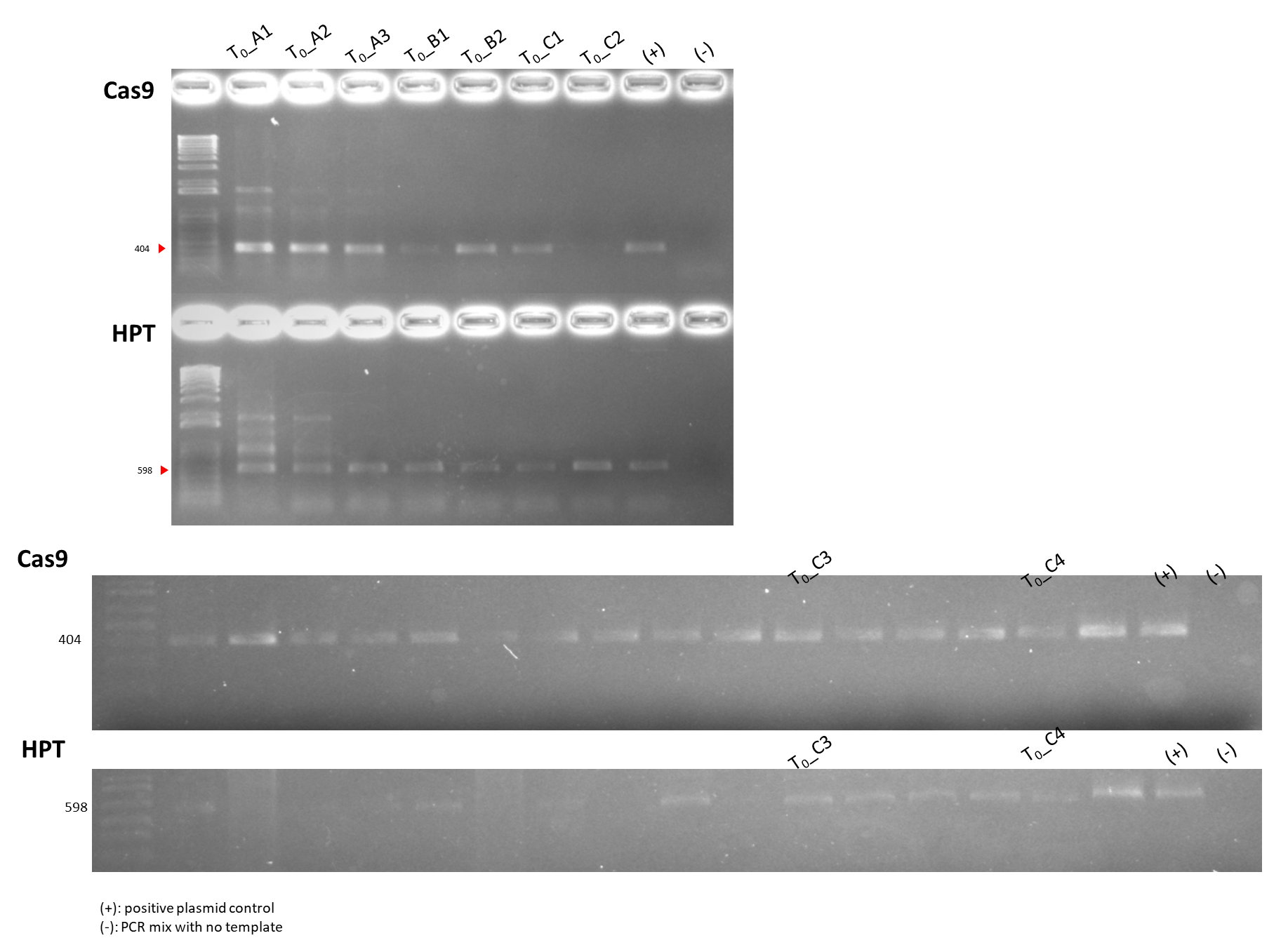

**Supplementary Figure 1.** Cas9 and HPT amplification from T_0_ plants.
